# Integration of strain and process optimization to increase autotrophic growth of engineered *Komagataella phaffii*

**DOI:** 10.1101/2025.05.21.655372

**Authors:** Michael Baumschabl, Lisa Lutz, Marina Jecmenica, Özge Ata, Diethard Mattanovich

## Abstract

Synthetic autotrophs are a promising platform for sustainable bioproduction using CO_2_ as substrate. The methylotrophic yeast *Komagataella phaffii* has been engineered to use CO_2_ as the sole carbon source by integration of the Calvin–Benson–Bassham (CBB) cycle, based on its native methanol assimilating xylulose monophosphate pathway (XuMP) cycle. Initial growth rates were low, but could be doubled by adaptive laboratory evolution (ALE). Beneficial mutations led to a decrease of CBB cycle reactions, indicating further limitations. During this study, temperature was identified as one of the key process parameters to improve autotrophic growth. For this reason, a new round of adaptive laboratory evolution was performed at the identified optimal cultivation temperature of 25°C, resulting in isolates growing up to 50 % faster compared to the control strain. Whole genome resequencing followed by reverse engineering helped to identify first key mutations of the evolved strains.

In addition, targeted engineering was performed by increasing the copy number of the key gene of the CBB cycle RuBisCO, which is the bottleneck of carbon fixation. Combining this with the optimal cultivation temperature boosted maximum specific growth rates of the autotrophic *K. phaffii* strain. In comparison to ALE, the targeted engineering still is lagging behind a bit. Starting from the initial condition, growth was boosted more than 2.5-fold in this study to a maximum of 0.025 h^-1^.

## Introduction

Nature has evolved CO_2_ assimilation several times, based both on linear and circular pathways. The most abundant pathway is the Calvin-Benson-Bassham (CBB) cycle. While energetically demanding it comes with the advantage that it works in aerobic conditions. The CBB cycle is described both in prokaryotic and eukaryotic organisms, *i*.*e*. in photosynthetic cyanobacteria as well as in chemolithoautotrophic bacteria (Hudson, 2024), and in the plant kingdom.

A few years ago, the integration of the CBB cycle to enable autotrophic growth of engineered, naturally heterotrophic microorganisms was described in *Escherichia coli* (*Gleizer et al*., *2019*) and the yeast *Komagataella phaffii* (previously known as *Pichia pastoris*) (Gassler et al., 2020). *K. phaffii*, equipped with a functional CBB cycle could grow directly in a CO_2_ rich atmosphere, while *E. coli* demanded initial adaptive evolution to become able to grow. Ben Nissan et al. demonstrated later that it requires three mutations to enable *E. coli* to grow on CO_2_ as the carbon source. Beside the beta subunit of the RNA polymerase (*rpoB*) and the cAMP receptor protein (*crp*), a mutation in phosphoglucoisomerase (*pgi*) directly interferes with a branching pathway of the CBB cycle (Ben Nissan et al., 2024).

To enhance growth on CO_2_ of engineered *K. phaffii* further, adaptive laboratory evolution was performed, leading to mutations in phosphoribulokinase (*PRK*), and in nicotinic acid mononucleotide adenylyltransferase (encoded by *NMA1*), slowing down their respective reaction rates. Further evolution led to mutations of *PEX5* encoding the peroxisomal receptor responsible for protein uptake into peroxisomes (Gassler et al., 2022) In addition, an increase of CO_2_ concentration (from 5 to 10%) (Baumschabl et al., 2022) and a decrease of oxygen supply (Baumschabl et al., 2023) proved to support faster CO_2_ assimilation.

Oxygen concentration is considered to be a critical parameter due to the oxygenation reaction of RuBisCO which deviates part of the carbon flux to 2-phosphoglycolate, wasting carbon and previously invested energy due to a decarboxylation step in photorespiration (Bauwe et al., 2010) which is, however, a necessary step to remove toxic 2-phosphglycolate. It is noteworthy that the synthetic autotrophic *K. phaffii* strain is using native enzymes for a similar pathway to remove the oxygenation byproduct which is probably one of the reasons for the efficient implementation of the CBB cycle in this yeast (Baumschabl et al., 2023).

The observed reduction of Prk activity by ALE points at an imbalance of the engineered CBB cycle. CBB is an autocatalytic cycle requiring a balance between the cycling metabolic flux and the output flux (Barenholz et al., 2017). Janasch et al. (2019)have demonstrated with a kinetic model of *Synechocystis sp*. That especially high concentrations of ribulose bisphosphate (the product of the Prk reaction) lead to unstable states of the CBB, supporting our observation that a rebalancing of Prk to adjust to the actual RuBisCO activity increases CBB flux. A decrease of oxygen concentration, however, alleviates the2ompetetion with CO_2_ for the RuBisCO active site so that higher CBB cycle rates are enabled without reduction of PRK activity. Lower cultivation temperatures may lead to a similar effect due to higher solubility of CO_2_ in the culture medium, and a more favourable relative dissolved CO_2_/O_2_ ratio at lower temperatures (Ku & Edwards, 1977).

It can be assumed that lowering temperature and oxygen concentration may lead to conditions where the benefit of the obtained Prk mutation is no longer exhibited, so that different growth optima may be reached with adaptive evolution of the parental CBB strain, potentially leading to higher specific growth rates. We have therefore investigated the effect of decreasing temperatures from 30°C to 25 °C and 20°C on specific growth rates of synthetic autotrophic *K. phaffii* strains, and performed ALE at lower temperatures. As an alternative strategy to balance the CBB cycle we increased the gene copy number of RuBisCO together with and without those of its chaperones GroEL and GroES.

## Results

### Lower temperature enhances growth of synthetic autotrophic *K. phaffii*

Strain engineering and optimization of the cultivation conditions are two key methods to improve growth of engineered microbial strains. Concerning culture conditions, multiple parameters including temperature and gas concentrations in the atmosphere can be fine-tuned to optimize growth of the synthetic autotrophic *K. phaffii* strain. We have previously shown that optimizing oxygen supply could improve growth rates significantly (Baumschabl et al., 2023). Based on the demonstrated impact of culture conditions we have studied the impact of temperature on specific growth rates.

Reducing the cultivation temperature from 30 °C to 25 °C boosted growth nearly 1.5-fold (Fig. 1), while further reduction to 20 °C had only a minor effect. Interestingly, the effect of temperature was slightly different in the reverse engineered version of the synthetic autotroph *K. phaffii* strain, which had a reduced activity of the Prk enzyme suppling the RuBisCO protein with its substrate ribulose 1,5-bisphosphate (Gassler et al., 2022). In this strain again reducing the temperature to 25 °C could increase growth significantly (Supplementary Fig 1). However, further reduction of the temperature decreased growth even below growth rates of the parental strain.

**Fig1:**
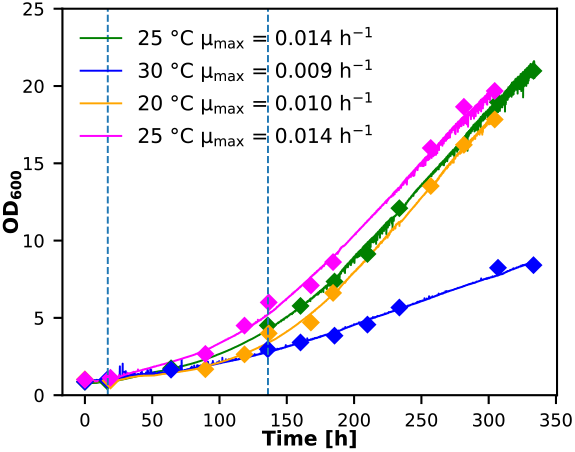
Bioreactor cultivations using different cultivation temperatures in the range of 20 °C and 30 °C to test their influence on growth. Diamonds: offline OD_600_ measurements; solid lines: online OD probe to monitor growth. Cultivations were performed at limited oxygen conditions and a CO_2_ concentration in the inlet air of 5%. Specific growth rates (µ) displayed in the figure were calculated in the interval indicated by the two dotted lines.

### Adaptive Laboratory Evolution finds new beneficial mutations at lower temperature

Adaptive laboratory evolution (ALE) proved to be a powerful tool to improve growth of the synthetic autotrophic *K. phaffii*. Previous ALE experiments were conducted at a cultivation temperature of 30 °C and could increase the growth rate 2-fold. However, cultivations of the evolved strains at reduced temperatures resulted in a weaker increase in growth at 25 °C, and at 20 °C even a reduction in growth in the strain harboring the mutation in the *PRK* gene compared to its parental strain. This was the motivation to perform a new round of ALE experiments starting again with the parental strain. The experiment was performed in triplicates at 25 °C for 266 days or 6379 hours of cultivation time. In addition, 3 cultures were exposed 4 times to UV light between 640 hours and 1900 hours of cultivation to increase mutation rate (denoted as EvoUV cultures). The course of the experiment is shown in Supplementary Fig 2 and Supplementary Fig 3. The dashed lines indicate the number of generations if growth would not have changed. After 4000 hours of cultivation time the number of generations increased over the theoretical number if no improvement had happened. In total the cultivations accumulated over 100 generations. Treating cells with UV slightly decreased the number of generations in the course of the evolution experiment because UV treatment decreased growth in the following passage of cells.

During the evolution experiment single cells were isolated three times after 108, 195 and 266 days by plating them on complex media to obtain single colonies. These were cultivated and growth was compared against the parental strain (Supplementary Fig. 3 A-C). Already after 108 days single colonies with significantly improved growth could be isolated. After the next isolation, clones with even higher growth rates could be isolated. The six best performing clones and the best from the first isolation (red lines in Supplementary Fig. 4 A, B) were selected and compared against the control and the engineered control (Fig 2 A). All clones grew faster than the control strain and even outperformed the engineered control. Maximum specific growth rate of the best clone was 0.023 h^-1^ in contrast to 0.016 h^-1^ for the control and 0.021 h^-1^ for the engineered control, respectively. Increasing the CO_2_ concentration to 10 % could further increase growth rates and again the isolated strains resulted in the highest growth rates compared to the control strains. These strains were also re-sequenced using Illumina whole genome sequencing to identify the mutations which occurred during ALE. Sequences were compared against the control strain, which was re-sequenced in parallel. To reduce complexity, only mutations which led to an amino acid change in proteins were further considered. In general, all tested single isolates showed at least 9 mutations which led to a change in an amino acid. Interestingly, the sequenced pools resulted in lower numbers of mutations compared to the single isolates. The data indicates that UV treatment resulted in a higher mutation rate. The sequencing results showed that there are only a few mutated genes which could be found in most isolates, namely *FLO11, PEX5, PIR1, SBT100*, PP7435_Chr3-1157, PP7435_Chr4-0629, PP7435_Chr4-1001 (Table 1).

**Table 1:** Overview of the sequencing results of the isolates. Number of mutations indicate the number of mutations which lead to an amino acid change. Parent pool indicates the pool the respective isolate originates from. A detailed overview of the amino acid changes of each gene can be found in the Supplementary Table 1.

| Strain | # Mutations | Parent pool | Genes Mutated |
| --- | --- | --- | --- |
| **EVO 8** | 11 | Evo pool II | *FLO11, PIR1, SBT100, PP7435_Ch3-1157, PP7435_Ch4-0629, PP7435_Ch4-1001* |
| **EVO2 10** | 9 | Evo pool II | *PP7435_Ch1-0080, PEX5, FLO11, PIR1, SBT100, PP7435_Ch3-1157, PP7435_Ch4-1001* |
| **EVO2 18** | 13 | Evo pool III | *PEX5, FLO11, PIR1, SBT100, PP7435_Ch3-1157, PP7435_Ch4-0629, PP7435_Ch4-1001* |
| **EVO2 19** | 9 | Evo pool III | *PEX5, FLO11, PIR1, SBT100, PP7435_Ch3-1157, PP7435_Ch4-0629, PpTRM1* |
| **EVO2 25** | 26 | Evo pool UV III | *RGT1, MTH1, ADH7, PEX5, FLO11, TAF2, PIR1, NSI1, TGL4, RSP5, MGM1, SBT100, PP7435_ch2-2727, RBG2, GCN2, PP7435_Ch3-0964, PP7435_Ch3-1157, SKG3, PP7435_Ch4-0629, MDS3, SEC16, ASF1, PP7435_Ch4-1001* |
| **EVO2 26** | 31 | Evo pool UV III | *PP7435_Ch1-0791, BPT1, DSS1, PP7435_Ch2-0160, PEX5, FLO11, TAF2, RIM9, PIR1, TRS130, PP7435_Ch2-0786, CUL3, SBT100, BUD7, GAD1, PP7435_Ch3-0835, PP7435_Ch3-1157, FLO5-2, EXO70, PP7435_Ch4-0629, TRC1, PP7435_Ch4-1001* |
| **EVO2 40** | 12 | Evo pool UV I | *UBP3, PEX5, FLO11, PIR1, ENV10, CBK1, PP7435_Ch3-1157, MHO1, DED1* |
| **EVO pool III** | 6 |  | *FLO11, PIR1, PP7435_Ch3-1157, PP7435_Ch4-0629* |
| **EVO UV pool III** | 8 |  | *PIR1, SBT100, ENV10, PP7435_Ch3-1157, PP7435_Ch4-1001* |

**Fig2:**
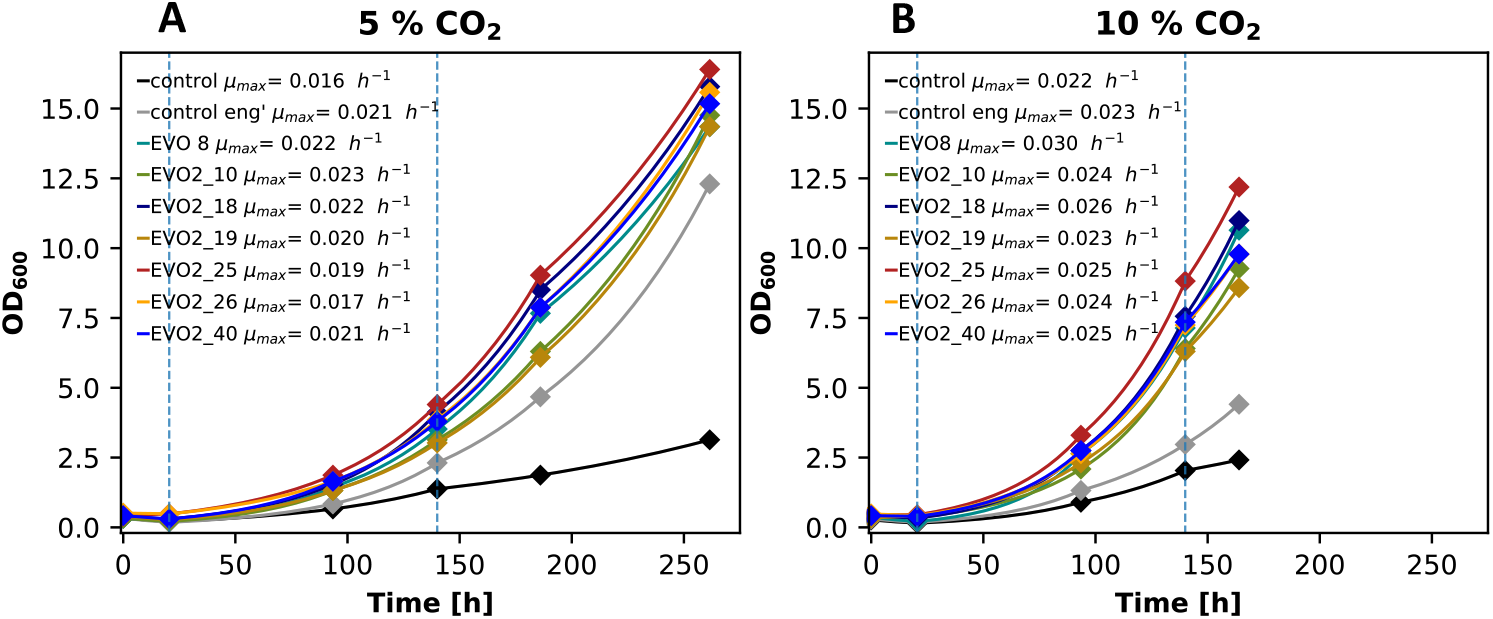
Growth curves of the evolution isolates cultivated at (A) 5% CO_2_ or (B) 10% CO_2_. Specific growth rates displayed in the figure were calculated in the interval indicated by the two dotted lines.

### Reverse engineering of selected mutations enhances autotrophic growth

Reverse engineering is a powerful tool to identify the mutations which are responsible for the improved phenotype. Two genes were considered most promising and were selected to implement single amino acid changing mutations in the parental strain (*PEX5* and *PIR1*). *PEX5* already appeared in earlier evolution experiments, also harboring a mutation at position 356 (but to a different amino acid). Both tested mutations of *PEX5* resulted in an increased specific growth rate up to 0.025 h^-1^, which was in a similar range as the evolution clones. In contrast, all tested mutations of *PIR1* did not result in a significant increase in growth compared to the control strain.

### Increasing RuBisCO expression supported faster autotrophic growth

Besides using ALE to increase growth we also employed rational engineering approaches to improve autotrophic growth rates. The rate limiting step of the CBB cycle is mostly considered to be the carboxylation reaction catayzed by RuBisCO. To boost CO_2_ fixation we tried to increase the copy number of the RuBisCO gene. In addition, in some clones the copy number of the *E. coli* chaperone genes *GroEL* and *GroES* were increased as well. This was done by integration of constructs harbouring a RuBisCO expression cassette (R clones), constructs including RuBisCO and the *E. coli* chaperone genes *GroEL* and *GroES* expression cassettes (RC clones) and a combination of both constructs to vary the ratio between RuBisCO and the chaperone copy numbers (RRC clones). Cultivations on 30 °C only showed minor differences in growth of the multi copy clones compared to the control strain. Highest maximum growth rates were achieved with strains harbouring 3 to 5 copies of the RuBisCO gene (Fig. 4). Based on our data a parallel increase of chaperone copy number did not lead to a significant increase in growth.

**Figure 3:**
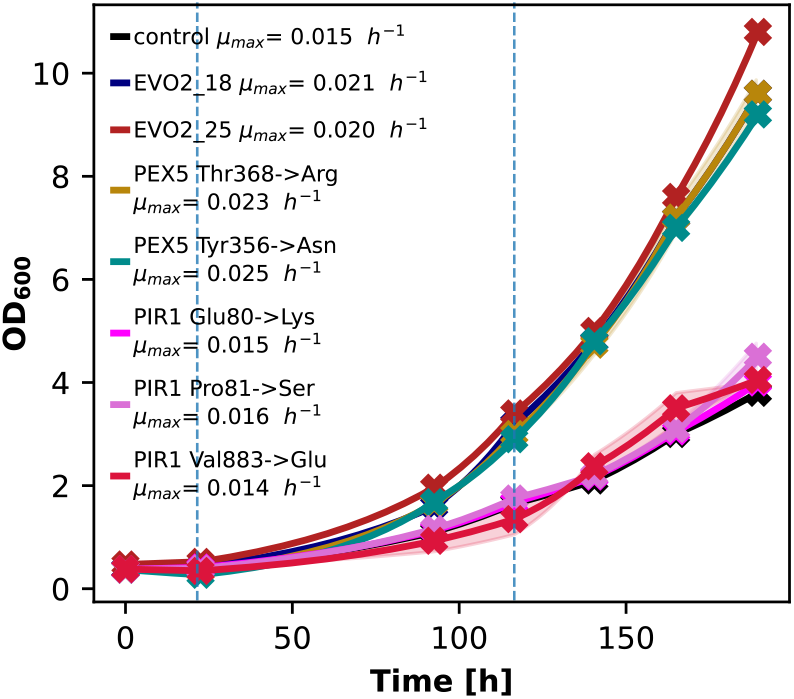
Results of the reverse engineering of selected mutations in PEX5 and PIR1. Shades indicate the standard deviation of the biological replicates. Specific growth rates displayed in the figure were calculated in the interval indicated by the two dotted lines.

**Figure 4:**
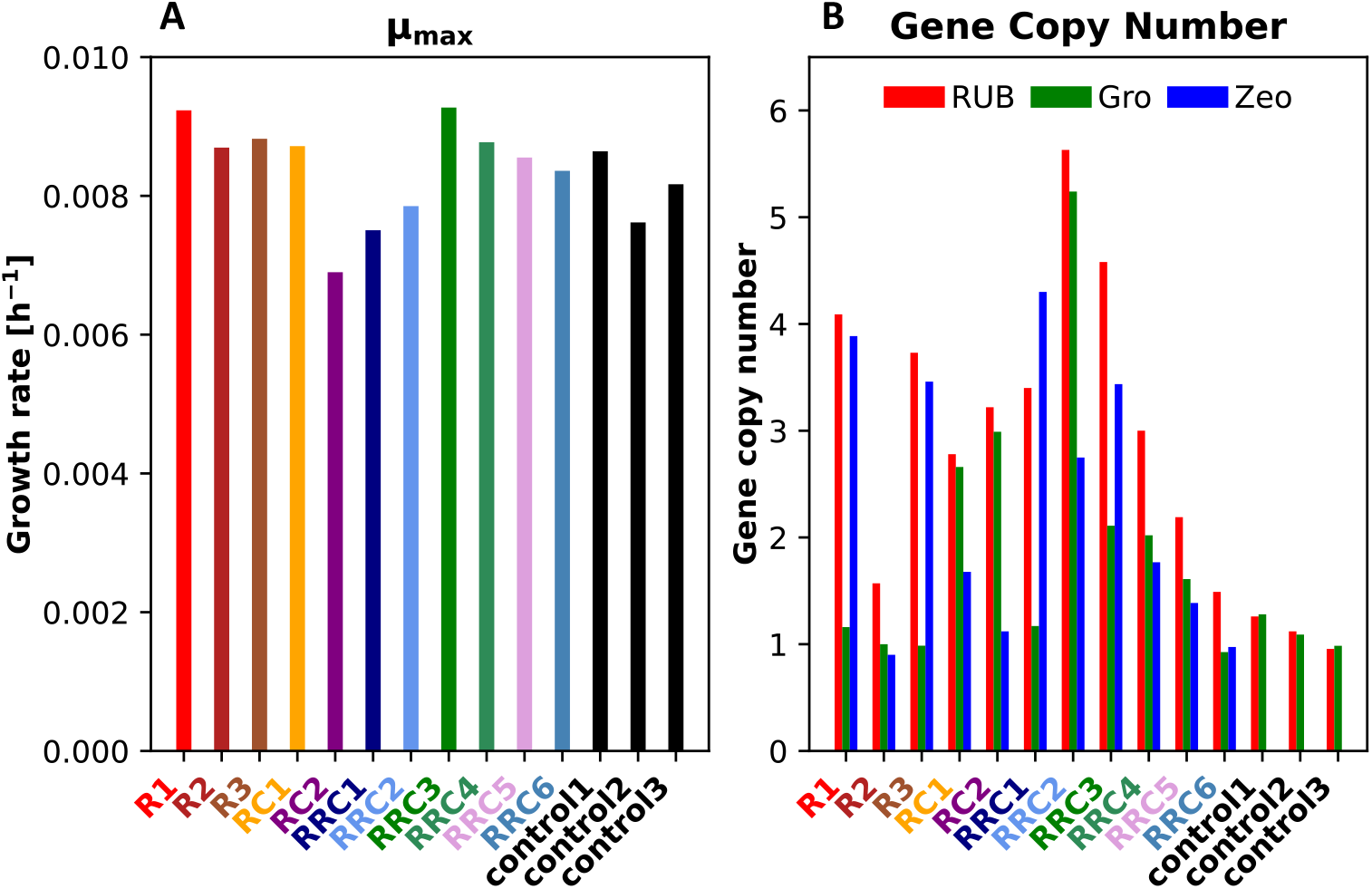
(A) Growth rates (calculated from 0 h to 140 h) of the multicopy clones compared against the single copy control strain cultivated at 30 °C. (B) Results of the gene copy number determination of the RuBisCO gene (RUB), the small subunit of the chaperones groES (Gro) and the zeocin resistance cassette (Zeo).

Since the clones were obtained by classical antibiotic selection, they also contained a high copy number of the antibiotic selection marker gene, mediating resistance to zeocin in this study. To remove the marker gene the Cre/Lox system was used. Unfortunately, this led to the loss of copy numbers of RuBisCO and chaperone genes, if the clones had more than 2 copies of the genes present. The loss in copies led to a reduction in maximum growth rate for most of the clones tested (Fig. 5).

**Figure 5:**
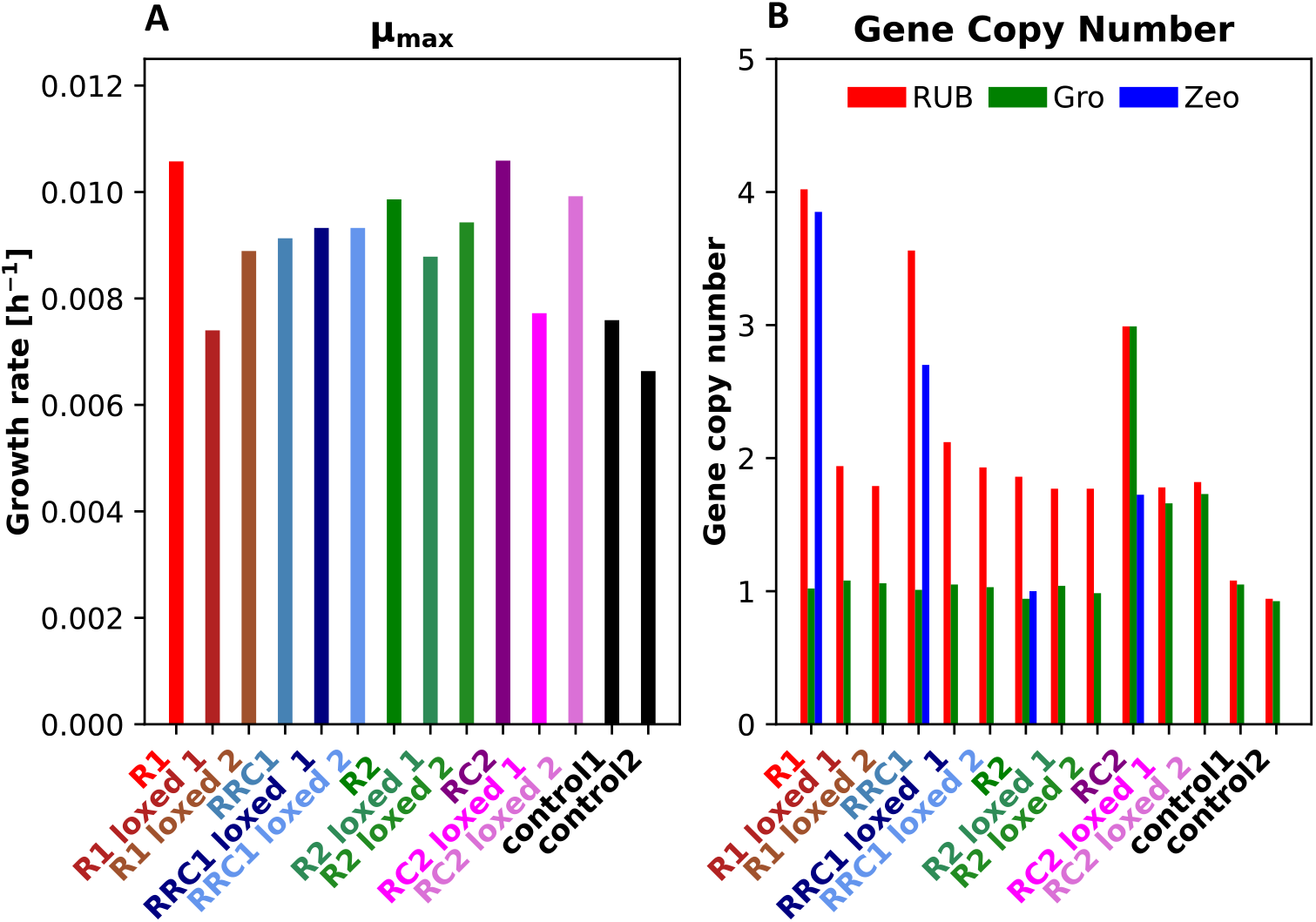
(A) Specific growth rates (calculated from 0 h to 70 h) of the multicopy clones and their respective loxed clones compared against the single copy control strain cultivated at 30 °C. (B) Results of the gene copy number determination of the RuBisCO gene (RUB), the small subunit of the chaperones groES (Gro) and the zeocin resistance cassette (Zeo).

Cultivation of the control strain at 25 °C compared to 30 °C increased the growth rate approximately 25%. Therefore, the effect of the temperature reduction was tested on selected clones (including their respective loxed clones). Surprisingly, reducing the temperature to 25 °C led much higher improvement in growth compared to cultivation at 30 °C. The best performing clone was RC2 with a maximum growth rate of 0.021 h^-1^. Cultivations at 25 °C showed a higher reduction in growth rate for the loxed clones which had a reduced number of copies (Fig. 6).

**Figure 6:**
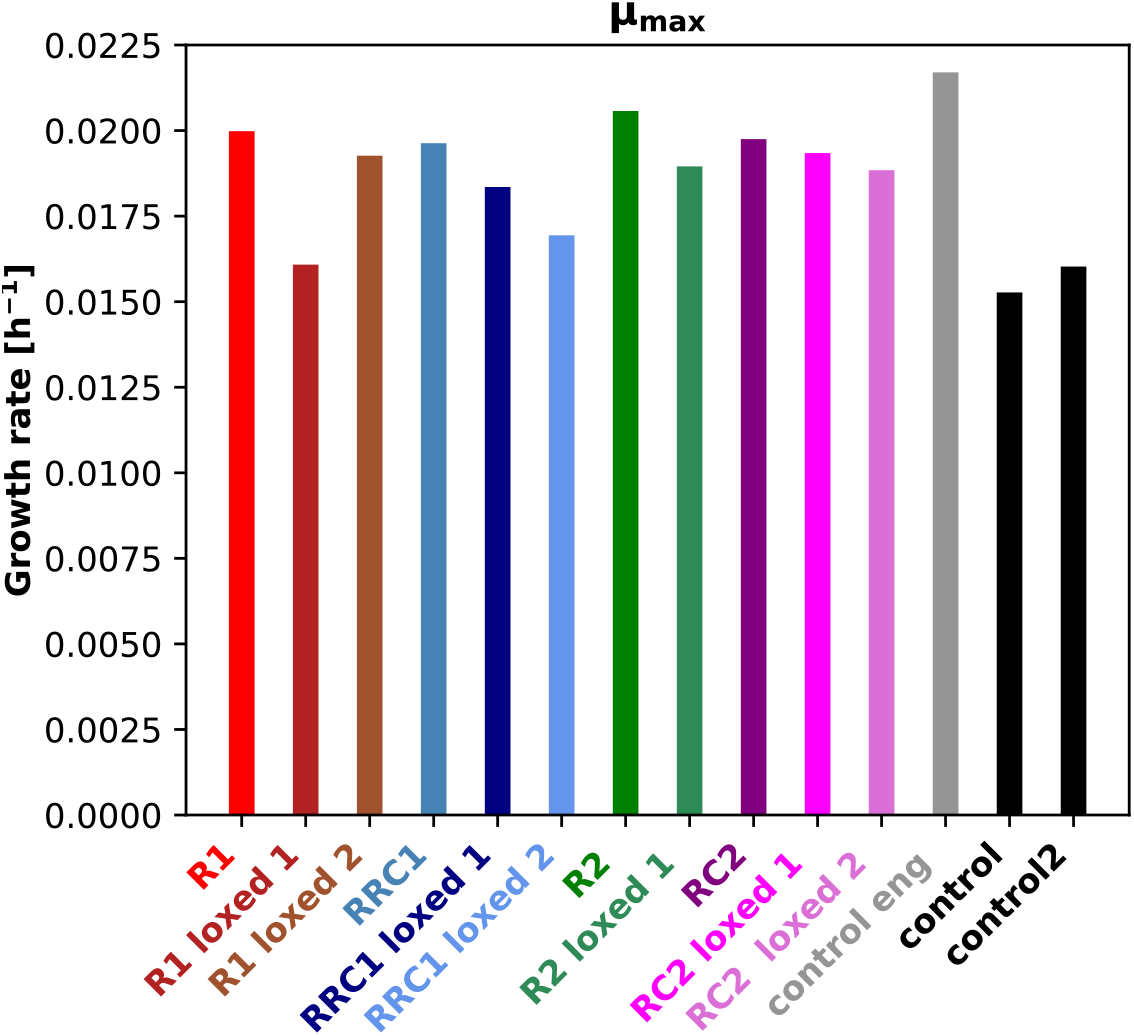
Specific growth rates (calculated from 24 h to 139 h) of the multicopy clones and selected loxed clones compared against the single copy control strain and the engineered control strain harboring a mutation in the PRK gene cultivated at 25 °C.

For the verification of the effect of the increased copy number of RuBisCO and chaperone genes, transcript levels of both were checked in cultivations at 25 °C. In addition, selected clones from the evolution experiment were tested as well. In all multicopy strains RuBisCO transcript levels were significantly increased. In addition, all clones had increased transcript levels of the chaperone gene, except RRC1 loxed 1, which only had a chaperone gene copy number of one. All evolution clones that were tested, however, resulted in similar transcript levels of both genes compared to the control strain. Both the multicopy clones and evolved clones grew faster than the parental strain (Fig. 7).

**Figure 7:**
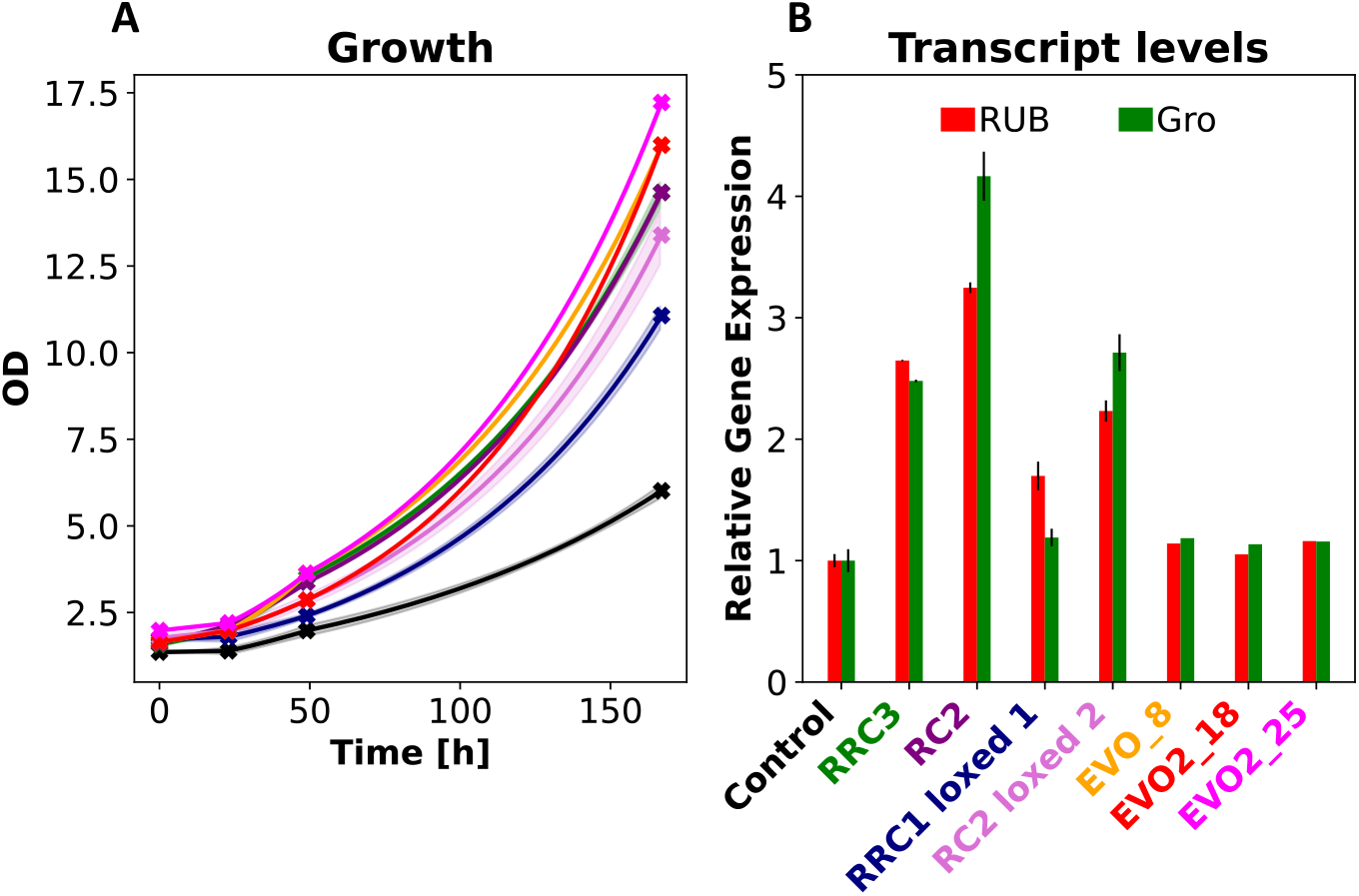
(A) Growth curves of the cultures performed for RNA extraction and transcript level determination. Shades indicate the standard deviation. (B) Gene expression levels of the RuBisCO gene (RUB) and the small subunit of the chaperones groES (Gro), relative to the control strain. Error bars indicate the standard deviation of the 3 biological replicates (EVO clones were performed as single experiments).

Finally, two selected high copy clones and one evolution isolate were compared using bioreactor cultivations at 25 °C as the cultivation temperature. As expected, all tested clones resulted in increased growth. Similar as in the shake flask cultivations, the evolution isolate grew slightly better compared to the high copy clones (Fig. 8).

**Figure 8:**
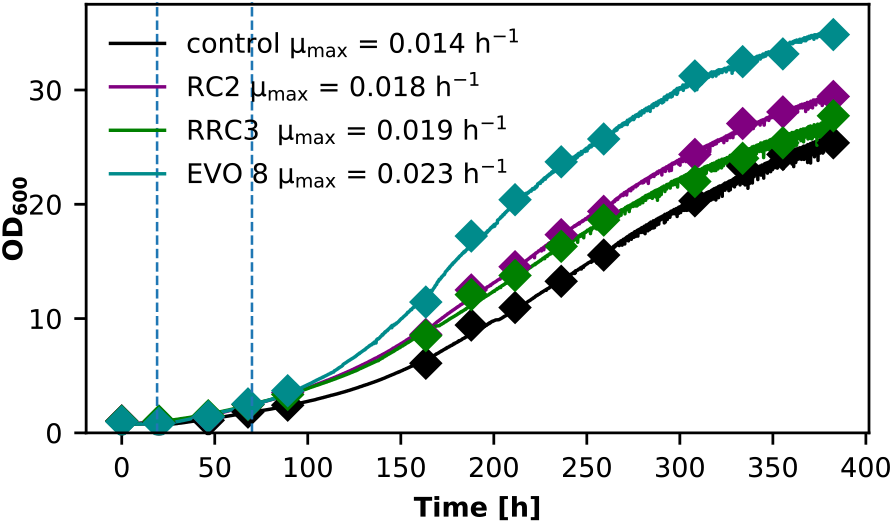
Bioreactor cultivations comparing the different engineering strategies against the control strain. Diamonds: offline OD_600_ measurements; solid lines: online OD probe to monitor growth. Cultivations were performed at limited oxygen conditions and a CO_2_ concentration in the inlet air of 5% Specific growth rates displayed in the legend were calculated in the interval indicated by the two dotted lines.

## Discussion

Integration of the CBB cycle enabled *K. phaffii* to use CO_2_ as its sole carbon source, however at rather slow growth rates. Growth could be enhanced by evolution, whereby mutations leading to a reduction of PRK activity had the best effect. Lower PRK activity likely alleviates an imbalance of the engineered CBB cycle (Gassler et al., 2020, 2022). Reduction of the available oxygen helped to improve growth as well. As a further process parameter, the effect of cultivation temperature was evaluated during this study. The optimal growth temperature of *K. phaffii* is 30 °C (Joseph et al., 2022). However, lowering the temperature was already reported beneficial for improving heterologous protein production in specific cases (Dragosits et al., 2009; Jahic et al., 2003; Li et al., 2001). For the synthetic autotrophic *K. phaffii* strain we could show that 25 °C is the optimal growth temperature, increasing the maximum growth rate roughly 50 percent compared to 30 °C. Interestingly, the effect of temperature was much lower for the reverse engineered strain which only differs by one point mutation in the *PRK* gene. Especially cultivation at 20 °C resulted in the highest differences in growth between the two tested strains. Whereas it resulted in a similar performance as at 25 °C in the parental strain, it led to the lowest growth in the engineered strain, even lower than the parental strain at 30 °C. We assume that the reduced activity of Prk, which leads to a balancing of the CBB cycle at 30 °C becomes a bottleneck at lower temperatures where more favorable CO_2_ availability and CO_2_/O_2_ ratio as well as better specificity of RuBisCO at lower temperatures remove the limits of the RuBisCO reaction (Gurrieri et al., 2019; Jordan & Ogren, 1984; Ku & Edwards, 1977).

Previous studies showed already that evolution of the synthetic autotrophic *K. phaffii* strains is a powerful tool to improve growth (Gassler et al., 2020, 2022). However, growth experiments at lower temperatures indicated in hindsight that the evolved clones were optimized to sub-optimal conditions. For this reason, a new round of evolution was started at a temperature of 25 °C. Again, mutants resulting in improved growth could be isolated. The best growing isolate outperformed the engineered control strain which so far resulted in the fasted autotrophic growth rates of *K. phaffii*. Increasing the temperature to 30 °C as used before still improves the growth rate of the isolate compared to the control, however the engineered control strain resulted in very similar growth. An increase of CO_2_ levels helped as well to further increase the gap of growth rates between the isolates and both control strains. This is a further indication that the mutation of Prk in the engineered control strain led to a reduction of the CBB cycle activity.

Since the ALE was performed longer compared to the previous attempts, more mutations which led to amino acid changes in proteins accumulated. The high number of mutated genes makes the evaluation of the crucial genes more difficult. Treatment with UV-light clearly increased the number of mutations, however no clear benefit in the final growth rates of the isolates could be observed. The highest number of mutations were found in *PIR1*, a gene encoding for a protein responsible in cell wall stability (Pal Khasa et al., 2011). Interestingly reverse engineering of a single mutation could not restore the improved growth phenotype. Another mutated gene which could be found in nearly every isolate is *PEX5*, a receptor for the peroxisomal targeting signal (Terlecky et al., 1995). Both tested single point mutation could nearly restore growth rates of the isolates indicating the importance of peroxisomal import for growth of the autotrophic strain. This is even more highlighted since mutations in *PEX5* (also in similar positions) were already occurring in previous evolution experiments, where it was shown that the mutation led to an increased import of proteins into the peroxisomes (Gassler et al., 2022). In addition, most of the isolates harbored mutations in the genes related to flocculation, *FLO11* and PP7435_Chr4-0629. Accumulation of mutations were also found in the two genes PP7435_Chr3-1157 and PP7435_Chr4-1001, both encoding hypothetical proteins, where functions are not known so far. The reverse engineering of this and previous work (Gassler et al., 2022) showed that ALE typically led to some crucial mutations which are responsible for the major part of the improved growth.

RuBisCO is known to be a bottleneck in carbon fixation by the CBB cycle (Prywes et al., 2025). Tests of variants with faster turn-overs (Davidi et al., 2020) did not lead to improvements in maximum growth rates in the autotrophic *K. phaffii* strain (Baumschabl et al., 2023). Therefore we attempted to increase the expression of the so far used RuBisCO by increasing the gene copy number, with and without that of the bacterial chaperones supporting RuBisCO folding. The effect on increased RuBisCO expression was negligible when cells were cultivated at 30 °C. A reason for this could be the lower CO_2_ solubility in the culture medium and a unfavorable CO_2_/O_2_ ratio, so that the cells cannot profit from the additional RuBisCO proteins. With a reduction of cultivation temperature these parameters change favorably so that the additional RuBisCO expression supported an increase of growth by 40 %, reaching similar growth rates compared to the reversed engineered clone which resulted in the fastest growth so far. A similar increase in growth rates after overexpression of the RuBisCO protein was already described in the autotrophic bacterium *Cupriavidus necator* (Kim et al., 2022) However in plants overexpression of RuBisCO on its own, did not lead to increased activity (Salesse-Smith et al., 2018; Suzuki et al., 2007). Between the different clones tested, differences in the transcript levels were observed. The transcript levels scale well with copy number for the chaperone gene, while for RuBisCO the transcript levels of RC2 were slightly higher compared to RRC3 even though it has less copies of the gene. However, this small differences in transcription did not result in big differences in growth. The removal of the resistance cassette using the Cre/Lox system reduced the copy number of RuBisCO and chaperones, which in the end resulted in a reduction of both, transcript levels and growth.

We have compared three approaches to increase specific growth rates of engineered autotrophic yeast strains. Optimization of cultivation temperature to 25 °C enhanced growth likely by an increase of CO_2_ solubility and the soluble CO_2_/O_2_ ratio, and it obviated that previous evolution at 30 °C was trapped in a local maximum, where mutations of PRK rebalanced the CBB cycle, however reducing its capacity at lower temperatures. Therefore, ALE was repeated at 25 °C, parallel to rational cell engineering by an increase of gene copy numbers of RuBiCO and its chaperones. Direct comparison of the two cell engineering approaches showed that ALE still outperforms rational engineering. The highest specific growth rate achieved in bioreactors was 0.023 h^-1^, which is close to methanol utilization slow (Mut^S^) strains of *K. phaffii*, which are often used in industry (Krainer et al., 2012; Looser et al., 2014).

## Material and Methods

### Strain generation

The initial strain used in this study was the synthetic autotrophic *K. phaffii* strain (CBBp + RuBisCO) published in (Gassler et al., 2020). The sequences of RuBisCO and the *E. coli* chaperones groEL and groES were taken from (Gassler et al., 2020). Plasmid were created using the Golden *Pi*CS system. (Prielhofer et al., 2017). The plasmids for increased gene copy number were linearized using *Asc*I (NEB) and 7 µg were transformed into electro-competent cells (Wu & Letchworth, 2004) and plated on YPD (10 g L ^-1^ yeast extract, 20 g L^-1^ soy peptone, 20 g L^-1^ glucose) plates containing 500 mg L^-1^ zeocin.

Reverse engineering was performed using the CRISPR-Cas9 system (Gassler et al., 2019). A gRNA cutting in the gene of interest was designed and donor DNA containing the desired point mutations was created. Positive clones were verified by Sanger sequencing of the desired region of the gene.

### Screening cultivation

Shake flask cultivations were performed according to Baumschabl et al., 2023. In brief, precultures were performed in YP media (10 g L ^-1^ yeast extract, 20 g L^-1^ soy peptone) containing 10 g L^-1^ glycerol over night at 25 °C and 180 rpm. Afterwards cells were washed once with water and used to inoculate main cultures with an OD of 0.5 or 1 using buffered Yeast Nitrogen Base media without amino acids (YNB) (3.4 g L^−1^, 100 mM potassium phosphate buffer pH 6, 10 g L^−1^ (NH_4_)_2_SO_4_ as nitrogen source and 0.5% (vol/vol) methanol at the beginning as energy source). Cultures were incubated either at 25 °C or 30 °C, 5 % CO_2_ and 180 rpm. After the first day the methanol concentration was increased to 1% (vol/vol). The cultivations were sampled every 2 to 3 days measuring optical density and methanol concentrations, which have been afterwards readjusted to 1% (vol/vol). Prior to every sampling, the evaporated volume was corrected using water.

### Evolution

The adaptive laboratory evolution was performed as follows: After a pre-culture in YP glycerol a 30 mL main culture in YNB was started in a 300 mL Erlenmeyer flask. Samples were taken regularly to check for optical density and methanol concentration which have been readjusted afterwards to 1% (vol/vol). After cultivations reached an optical density of 15-20, cryos of each pool were prepared and cells were diluted again to an optical density of 1, or later 0.5. For the EVO UV pools freshly diluted cells were exposed to UV light by putting the shake flasks on a UV transilluminator (TFP-M/WL, MWG Biotech) after 26, 34, 66 and 80 days for 1 to 5 minutes. After 108, 195 and 266 days of adaptive laboratory evolution single clones were isolated by diluting the cultures followed by plating them YP glucose (20 g L^-1^).

### Gene copy number determination

The gene copy number determination was performed according to Staudacher et al., 2022. Cells were cultivated overnight in YP media. Genomic DNA was isolated using the Wizard^®^ Genomic DNA Purification Kit (Promega). Three µL of the genomic DNA dilutions with a DNA concentration of 2.7 ng µL^-1^ were used as template for the qPCR runs, which were performed using the 2x qPCR S’Green BlueMix (Biozym) and primers binding in the RuBisCO gene, the small subunit of the *E. coli* chaperone *groES*, the zeocine resistance cassette and *K. phaffii ACT1* used as housekeeping gene. All samples were analyzed using 4 replicates. Quantification was performed using the comparative quantification method of the Rotorgene software (Qiagen).

### Determination of transcript levels

The extraction of RNA and following transcript level determination was performed according to Heistinger et al., 2018. An aliquot of 10 OD_600_ units of the screening culture was harvested by centrifugation after 48 hours of cultivation. The pellet was resuspended in 1 mL TRI reagent solution (Invitrogen) and the RNA was isolated according to the protocol. Remaining DNA was removed using the DNA-free™-kit (Invitrogen). cDNA was synthesized using the cDNA Synthesis Kit (Biozym) and the Oligo d(T)23 VN primer (NEB).

The qPCR runs were performed using the 2x qPCR S’Green BlueMix (Biozym) and primers binding in the RuBisCO gene, the small subunit of the *E. coli* chaperone *groES*, the zeocine resistance cassette and *K. phaffii ACT1* used as housekeeping gene. All samples were analyzed using 4 replicates. Quantification was performed using the comparative quantification method of the Rotorgene software (Qiagen).

### Genomic DNA extraction and Next Generation Sequencing

Genomic DNA of the parental strain, the evolved strains and the pooled samples was extracted using the Qiagen Genomic-tip 20/G kit (Qiagen) according to manufacturer’s recommendations. The strains were grown in YPD medium (10 g L^-1^ yeast extract, 20 g L^-1^ soy peptone, 2% glucose) until an OD_600_ of 10 was reached and the appropriate cell number was harvested by centrifugation.

Samples were sent to the Next Generation Sequencing Facility at the Vienna BioCenter Core Facilities (VBCF), which are part of the Vienna BioCenter (VBC), where library preparation, sequencing and demultiplexing was performed. Sequencing was done on an Illumina NextSeq2000 P1 (PE300) platform to a coverage of > 30X for single strains and > 100X for pooled samples, respectively.

All further steps were carried out as described in Gassler et al. 2022. In brief, reads were analyzed using CLC Genomic Workbench 12.0 (Qiagen), where they were first mapped against the reference genome of CBS7435 (Valli et al., 2016). Basic variant detection was performed setting the ploidy parameter to 1, and obtained variants were filtered against those detected within the parental strain for each evolved strain/pool separately. All remaining variants were compared across all evolved samples and most likely candidates were considered for reverse engineering.

### Bioreactor cultivation

Bioreactor cultivations were performed similarly to Baumschabl et al., 2023, using a DASGIP system (Eppendorf) using YNB media, (NH_4_)_2_SO_4_ as nitrogen source, 10 mM phosphate buffer pH 6 and 0.5 % (vol/vol) methanol at the start. Oxygen in the inlet gas was limited, so that the dissolved oxygen concentration was not exceeding 15%. The pH was regulated by using potassium hydroxide as base.

Bioreactors were inoculated with an OD_600_ of 1 grown in YP glycerol (10 g L^-1^) overnight. Cells were washed with water twice before inoculation. After inoculation the first sample was taken and on the next day and the methanol concentration was increased to 1 % (vol/vol).

### HPLC measurements

Methanol concentrations were quantified by HPLC using the method published in (Baumschabl et al., 2022).

## Acknowledgements

The COMET center acib: Next Generation Bioproduction is funded by BMIMI, BMWET, SFG, Standortagentur Tirol, Government of Lower Austria and Vienna Business Agency in the framework of COMET – Competence Centers for Excellent Technologies. The COMET Funding Program is managed by the Austrian Research Promotion Agency FFG. MB and DM were supported by the Austrian Science Fund (Grant-DOI 10.55776/W1224, Doctoral Program on Biomolecular Technology of Proteins (BioToP)).

## Author contributions

ÖA and DM designed research, MB, LL, and MJ performed the experiments, MB, MJ analyzed the data, MB, ÖA and DM drafted the manuscript. All authors read and approved the manuscript.

