## Supplemental figure for "Integration of strain and process optimization to increase autotrophic growth of engineered *Komagataella phaffii*"

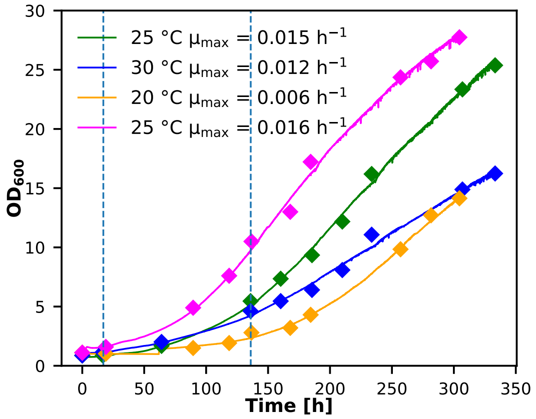


Supplementary Figure 1: Bioreactor cultivations of the reverse engineered strain habouring a mutation in the *PRK* gene using different cultivation temperatures in the range of 20 °C and 30 °C to test their influence on growth. Diamonds: offline $OD_{600}$ measurements; solid lines: online OD probe to monitor growth. Cultivations were performed at limited oxygen conditions and a $CO_2$ concentration in the inlet air of 5%. Specific growth rates displayed in the figure were calculated in the interval indicated by the two dotted lines.

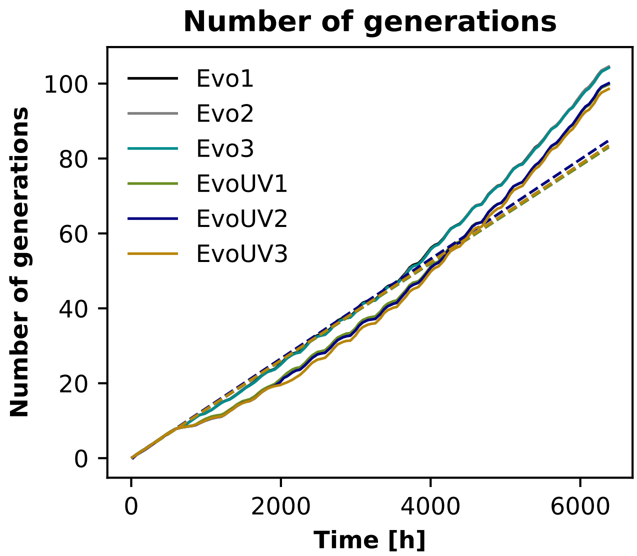


Supplementary Figure 2: Accumulation of number of generations of the different ALE cultivations during the course of the ALE experiment. The dashed lines indicate the theoretical numbers of generations if the growth rate would not have been improved.

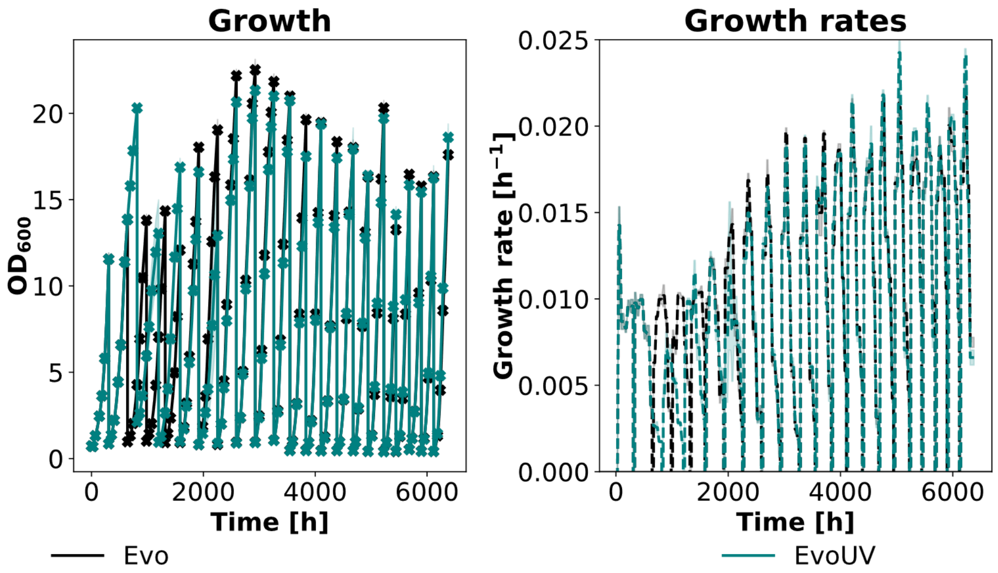


Supplementary Figure 3: Time course of the ALE experiment. The average OD and specific growth rates of all non treated cultures (black lines) and UV treated cultures (green lines) is shown.

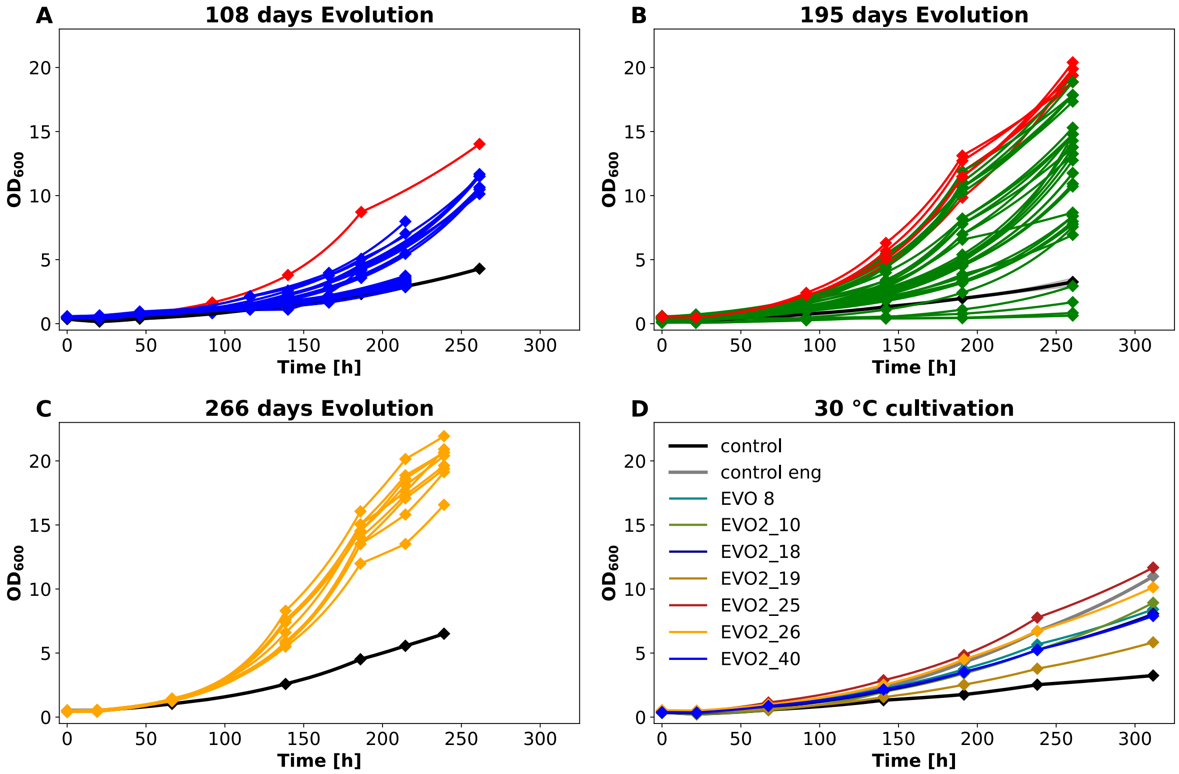


Supplementary Figure 4: Growth at 25 °C of isolates of the ALE experiment isolated after (A) 108 days, (B) 195 days or (C) 266 days of cultivation. The black line indicates the parental strain, and the red curves indicate the isolates which where further selected for whole genome re-sequencing. (D) Growth curves of the evolution isolates cultivated at 30 °C. The black line indicates the parental strain and the grey line the reversed engineered strain harboring a mutation in the *PRK* gene.

Supplementary Tabel 1: Overview of the whole genome re-sequencing results

| | | Strain | | | | | | | | |
|---|---|---|---|---|---|---|---|---|---|---|
| **Gene** | **Mutation** | EVO 8 | EVO2 10 | EVO2 18 | EVO2 19 | EVO2 25 | EVO2 26 | EVO2 40 | EVO pool III | EVO UV pool III |
| | | Evo pool II | Evo pool II | Evo pool III | Evo pool III | Evo pool UV III | Evo pool UV III | Evo pool UV I | | |
| **Number of Mutations** | | 11 | 9 | 13 | 9 | 26 | 31 | 12 | 6 | 8 |
| **Flo11** | Asn40Asp | x | x | x | x | x | x | x | x | |
| **PP7435_Chr1-0080** | Ser392f | | x | | | | | | | |
| **RGT1** | Pro79Ser | | | | | x | | | | |
| **MTH** | Ser87Phe | | | | | x | | | | |
| **ADH7** | Gly58Ser | | | | | x | | | | |
| **PP7435_Chr1-0791** | Leu19Phe | | | | | | x | | | |
| **BPT1** | Ser413Phe | | | | | | x | | | |
| **DSS1** | Leu909Phe | | | | | | x | | | |
| **PP7435_Chr2-0160** | Pro346Leu | | | | | | x | | | |
| **UBP3** | Thr464Ile | | | | | | | x | | |
| **PEX5** | Asn272Ile | | x | | | | | | | |
| | Thr368Arg | | | x | | | | | | |
| | Ala364_Ala365insGlyTyrAspAsnAla | | | | x | | | | | |
| | Tyr356Asn | | | | | x | | | | |
| | Tyr356Cys | | | | | | x | | | |
| | Asn363_Ala364insAspGlyTyrAspAsn | | | | | | | x | | |
| **PIR1** | Glu80Lys | x | | x | x | x | x | | | |
| | Pro81Ser | x | x | x | x | x | x | x | x | x |
| | Val83Glu | x | x | x | x | x | x | x | x | x |
| | Val131Ser | x | | | | | | | | |
| | Gln133Glu | x | | | | | | | x | |

| | | | | | | | | | | |
|---|---|---|---|---|---|---|---|---|---|---|
| | Thr135Ala | x | | | | | | | | |
| **SBT100** | Gln1312His | x | x | x | x | x | x | | | x |
| **PP7435_Chr3-1157** | Leu245Ser | x | x | x | x | x | x | x | x | x |
| | Ser290Gly | | | x | | | x | | | |
| | Ser288Trp | | | x | | | x | | | |
| | Tyr281fs | | | x | | | x | x | | |
| | Val279fs | | | x | | x | x | x | | |
| | Asp293Glu | | | | | | x | | | |
| | Ile287Pro | | | | | | x | | | |
| **PP7435_Chr4-0629** | Lys157Gln | x | | x | x | x | x | x | x | |
| **PP7435_Chr4-1001** | Val34Thr | x | x | x | | x | x | | | x |
| | Lys36Glu | | x | | | | x | | | x |
| | Tyr39His | | | | | | | | | x |
| **TRM1** | Ser619fs | | | | x | | | | | |
| **TAF2** | Glu1227Lys | | | | | x | x | | | |
| **NSI1** | Ser25Phe | | | | | x | | | | |
| **TGL4** | Tyr98Phe | | | | | x | | | | |
| **RSP5** | Leu131Phe | | | | | x | | | | |
| **MGM1** | Glu278Lys | | | | | x | | | | |
| **PP7435_chr2-2727** | Ile287Lys | | | | | x | | | | |
| **RBG2** | Ser363Phe | | | | | x | | | | |
| **GCN2** | Gly1672Asn | | | | | x | | | | |
| **PP7435_Chr3-0964** | Gly915Glu | | | | | x | | | | |
| **SKG3** | Thr523Ile | | | | | x | | | | |
| **MDS3** | Gly495Ser | | | | | x | | | | |
| **SEC16** | Pro838Ser | | | | | x | | | | |
| **ASF1** | Lys17Glu | | | | | x | | | | |
| **RIM9** | Leu102Pro | | | | | | x | | | |

| | | | | | | | | | | |
|---|---|---|---|---|---|---|---|---|---|---|
| **TRS130** | Gly1074Glu | | | | | | x | | | |
| **PP7435_Chr2-0786** | Tyr26Phe | | | | | | x | | | |
| **CUL3** | Pro739Leu | | | | | | x | | | |
| **BUD7** | Ser245Phe | | | | | | x | | | |
| **GAD1** | Ser147Phe | | | | | | x | | | |
| **PP7435_Chr3-0835** | Val217Met | | | | | | x | | | |
| **FLO5-2** | Phe557Tyr | | | | | | x | | | |
| **EXO70** | Ser133Cys | | | | | | x | | | |
| **TRC1** | Leu266Phe | | | | | | x | | | |
| **CBK1** | Pro320Leu | | | | | | | x | | |
| **MHO1** | Tyr233His | | | | | | | x | | |
| **DED1** | Ile397Arg | | | | | | | x | | |
| **EVN10** | Leu210fs | | | | | | | | | x |
